# Learning a latent representation of human genomics using Avocado

**DOI:** 10.1101/2020.06.18.159756

**Authors:** Jacob Schreiber, William Noble

## Abstract

In the past decade, the use of high-throughput sequencing assays has allowed researchers to experimentally acquire thousands of functional measurements for each basepair in the human genome. Despite their value, these measurements are only a small fraction of the potential experiments that could be performed while also being too numerous to easily visualize or compute on. In a recent pair of publications, we address both of these challenges with a deep neural network tensor factorization method, Avocado, that compresses these measurements into dense, information-rich representations. We demonstrate that these learned representations can be used to impute with high accuracy the output of experimental assays that have not yet been performed and that machine learning models that leverage these representations outperform those trained directly on the functional measurements on a variety of genomics tasks. The code is publicly available at https://github.com/jmschrei/avocado.

---

The field of genomics is undergoing a surge in the number of high quality, publicly available data sets. These data sets include genome-wide measurements of several types of biochemical activity such as chromatin accessibility, transcription, protein binding, and histone modification. Understanding this biochemistry is crucial for explaining the molecular basis for cellular phenomena, such as aging and disease. As a result, large collaborative efforts such as the Roadmap Epigenomics Mapping Consortium (Kundaje et al., 2015) and the NIH ENCODE Project (ENCODE Project Consortium, 2012) have prioritized performing experiments that measure dozens of forms of functional activity in hundreds of human cell types, primary cell lines, and tissues (“biosamples”). Today, the ENCODE Compendium is one of the most comprehensive resources for genomics data sets in the world, with over 10,000 data sets available.

Unfortunately, there are two major problems with compendia such as ENCODE’s. First, these compendia are now massive in size, containing thousands of data sets that require hundreds of gigabytes to store after processing and compression. It would be difficult for a researcher to visualize even a small subset of these data sets at a particular region or to perform computation on all of them without appropriate hardware. Second, despite their size, these compendia are generally incomplete. For example, the ENCODE Compendium has fewer than 1% of the experiments that could potentially be performed. This sparsity poses difficulties for computational methods that rely on a common set of functional measurements in each biosample, or for investigators whose research happens to be on a biosample that has had very few experiments performed in it.

We address both these challenges with Avocado, a deep tensor factorization model. Avocado organizes a set of genomics experiments into a 3D tensor whose axes correspond to biosample, assay type, and genomic position (Fig 1A). We refer to the model as “deep” because the dot product operation in standard factorization methods is replaced with a deep neural network (Fig 1B). The latent representations and neural network weights are trained using standard gradient descent methods on the regression task of predicting values within the tensor.

**Figure 1.**
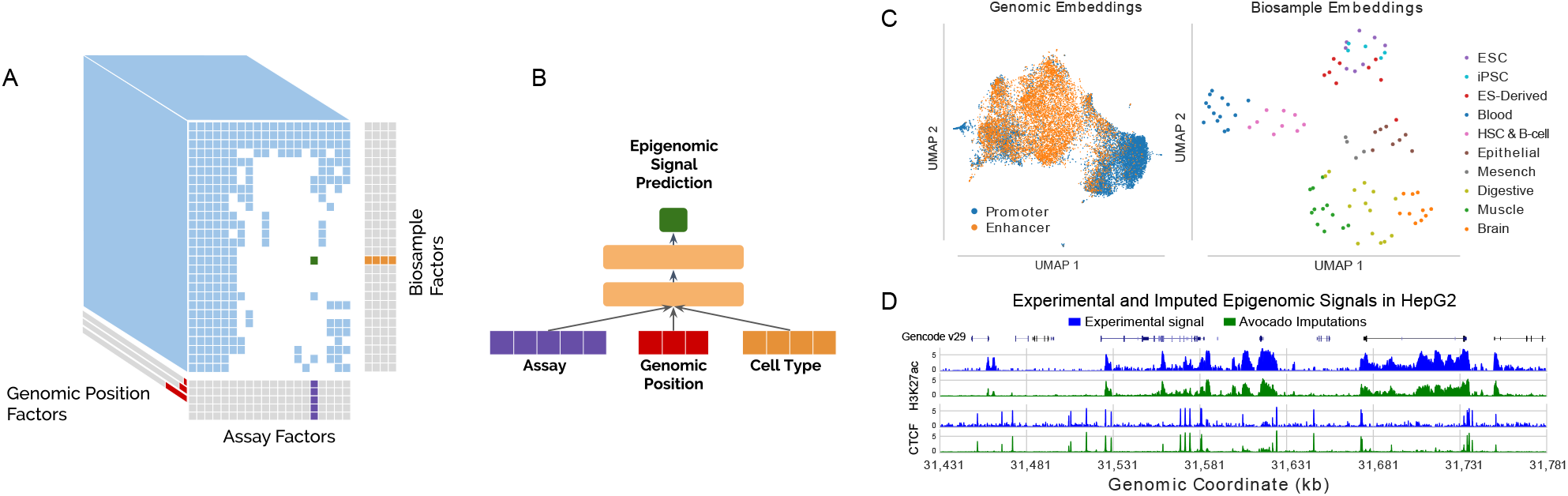
An overview of the Avocado model. (A) Genomics experiments are organized into a 3D tensor with the axes corresponding to biosample, assay, and genomic position, and Avocado learns latent representations for these three axes. (B) Avocado uses a neural network whose input is the concatenation of latent factors and whose output is the corresponding experimental signal. (C) UMAP projections of the genomic position and cell type representations with points colored by enhancer/promoter identity or anatomy type respectively. (D) Example experimental signal and imputations for the histone modification H3K27ac and binding of the protein CTCF at the same positions in the cell type HepG2. GENCODEv29 annotations indicate the locations of genes.

In a pair of publications, we apply Avocado to the Roadmap Compendium (Schreiber et al., 2020b) and then to the much larger ENCODE Compendium (Schreiber et al., 2020a). Our first main observation was that the learned latent representations encode complex biology. For example, a projection of the genome representations revealed a continuum between annotated enhancers and promoter elements, and a projection of the biosample representations showed a clustering by anatomy type (Fig 1C). Our second main observation was that the imputations made from the model were high quality and more accurate than previous approaches (Fig 1D). Additionally, we found that Avocado’s imputed transcription factor binding tracks outperformed the top participants in a recent ENCODE-DREAM transcription factor binding challenge (https://www.synapse.org/#!Synapse:syn6131484/wiki/402026). Overall, the imputations completing the ENCODE Compendium covered 400 human biosamples and 84 assays. This set of >30k genome-wide imputations represent, to our knowledge, the largest imputation of genomics experiments that has been performed to date, and the first time that this many forms of biochemistry were jointly modeled.

We anticipate that researchers will find Avocado’s latent representations to be widely useful. For instance, when used as input in the place of functional measurements, we found that these representations improved the performance of machine learning models trained to predict gene expression, promoter-enhancer interaction, replication timing, and frequently interacting regions (FIREs). Any model that currently takes functional measurements as input would likely benefit from instead using Avocado’s representations. Further, the representations can trivially be used to calculate a similarity between each pair of biosamples or each pair of genomic positions. These similarities provide a natural way to identify a functionally diverse set of biosamples that a new assay should be applied to or a functionally diverse set of genomic positions that expensive assays, such as tiling arrays, should profile.

We expect that the main value of the imputations will come from expanding the utility of existing computational methods and aiding researchers in hypothesis generation. Computational methods that rely on a common set of assays can substitute in imputations when experimental data is not available, allowing them to be comprehensively performed on all biosamples in the ENCODE Compendium. In cases where experimental data is available, we have observed that using the imputations instead can lead to improved model performance in part becase the imputations serve as a de-noised version of the experimental data. Additionally, imputations can be inspected to identify interesting patterns, such as clusters of biosamples exhibiting unexpected functional activity at a locus, that should be followed-up with experimental validation.

The code, models, and learned latent representations can be found at https://github.com/jmschrei/avocado. The >36k imputed genome-wide data sets produced during these projects can be found on the ENCODE portal https://www.encodeproject.org and are grouped by publication under the accessions ENCSR617ILB and ENCSR481OSA.

## Notes

### Competing Interest Statement

The authors have declared no competing interest.

